# Transcriptional atlas of the human immune response to 13 vaccines reveals a common predictor of vaccine-induced antibody responses

**DOI:** 10.1101/2022.04.20.488939

**Authors:** T. Hagan, B. Gerritsen, LE. Tomalin, S. Fourati, MP. Mulè, DG. Chawla, D. Rychkov, E. Henrich, HER. Miller, J. Diray-Arce, P. Dunn, A. Lee, The Human Immunology Project Consortium (HIPC), O. Levy, R. Gottardo, M. M. Sarwal, JS. Tsang, M. Suárez-Fariñas, RP. Sékaly, SH. Kleinstein, B. Pulendran

**Affiliations:** Division of Infectious Diseases, Cincinnati Children’s Hospital Medical Center, Cincinnati, OH, USA; Department of Pediatrics, University of Cincinnati College of Medicine, Cincinnati, OH, USA; Yale School of Medicine, New Haven, CT, USA; Center for Biostatistics, Icahn School of Medicine at Mount Sinai, New York, New York, USA; Emory University School of Medicine, Atlanta, GA, USA; Multiscale Systems Biology Section, Laboratory of Immune System Biology, NIAID and Center for Human Immunology (CHI), NIH, Bethesda, MD, USA; NIH-Oxford-Cambridge Scholars Program, Cambridge University, Cambridge, United Kingdom; Division of Transplant Surgery, University of California, San Francisco, San Francisco, CA, USA; Fred Hutchinson Cancer Research Center, Seattle, WA, USA; Precision Vaccines Program, Boston Children’s Hospital, Boston, MA, USA; Harvard Medical School, Boston, MA, USA; ImmPort Curation Team, NG Health Solutions, Rockville, MD, USA; Institute for Immunity, Transplantation, and Infection, Stanford University School of Medicine, Stanford University, Stanford, CA, USA; Broad Institute of MIT and Harvard, Cambridge, MA; University of Lausanne and Lausanne University Hospital, Lausanne, Switzerland; Swiss Institute of Bioinformatics, Lausanne, Switzerland; NIAID, NIH, Bethesda, MD, USA; Drexel University, Philadelphia, PA, USA; University of California, Berkeley, Berkeley, CA, USA; Columbia University Medical Center, New York, NY, USA; Boston Children’s Hospital, Harvard Medical School, Boston, MA, USA; La Jolla Institute for Immunology, La Jolla, CA, USA; Stanford University School of Medicine, Stanford University, Stanford, CA, USA; Icahn School of Medicine at Mount Sinai, New York, New York, USA; David Geffen School of Medicine at University of California, Los Angeles, CA, USA; University of California, San Francisco, San Francisco, CA, USA; Seattle Children’s Research Institute, Seattle, WA, USA; Laboratory of Immune System Biology, NIAID and NIH Center for Human Immunology (CHI), NIH, Bethesda, MD, USA

## Abstract

Systems biology approaches have been used to define molecular signatures and mechanisms of immunity to vaccination. However, most such studies have been done with single vaccines, and comparative analysis of the response to different vaccines is lacking. We integrated temporal transcriptional data of over 3,000 samples, obtained from 820 healthy adults across 28 studies of 13 different vaccines and analyzed vaccination-induced signatures associated with the antibody response. Most vaccines induced similar kinetics of shared transcriptional signatures, including signatures of innate immunity occurring 1-3 days post-vaccination, as well as the canonical plasmablast and cell cycle signatures appearing 7 days post-vaccination. However, the yellow fever vaccine YF-17D uniquely induced an early transient signature of T and B cell activation at Day 1, followed by delayed antiviral/interferon and plasmablast signatures that peaked at Days 7 and 14-21, respectively. Thus, despite the shared transcriptional response to most vaccines, at any given time point there was no evidence for a “universal signature” that could be used to predict the antibody response to all vaccines. However, accounting for the asynchronous nature of responses led to the identification of a time-adjusted signature that improved prediction antibody of responses across vaccines. These results provide a transcriptional atlas of the human immune response to vaccination and define a common, time-adjusted signature of antibody responses to vaccination.

## Introduction

Systems vaccinology employs high-throughput -omics measurements together with systems-based analysis approaches to better understand immune responses to vaccination^1, 2^. The recent growth of this field, which began with initial studies of yellow fever^3, 4^ and seasonal influenza^5, 6^ vaccines, has rapidly expanded to include studies profiling responses to a range of vaccines and vaccine platforms, including those targeting diverse pathogens and age groups^7-20^. These studies have led to important discoveries such as the role for the nutrient sensor general control nonderepressible 2 (GCN2) in enhancing antigen presentation during responses to yellow fever vaccination^21^, as well as the impact of the gut microbiota in promoting antibody responses to inactivated influenza vaccination^22, 23^. However, outside of a few studies^8-10^, thus far the vast majority have examined immune responses to a single vaccine, hindering the ability to contextualize the findings and understand how differences in vaccine formulations can impact immunogenicity.

Another important outcome of such studies has been the identification of early transcriptional signatures predictive of immune response quality such as subsequent antibody^3, 5, 18^ or antigen-specific T cell^3, 7, 14^ responses. These findings may enable more rapid and personalized evaluation of vaccine efficacy and development of improved next-generation vaccines. Yet again, a current limitation is that the identified predictive signatures thus far have been described in the context of responses to a single vaccine, and the extent to which predictors of immune response quality are conserved across vaccines is unclear^24, 25^. We previously sought to address the question of whether there was a ‘universal signature’ that could be used to predict antibody responses to any vaccine by analyzing the transcriptional response to 5 different human vaccines. Our analysis revealed distinct transcriptional signatures of antibody responses to different classes of vaccines, and provided key insights into primary viral, protein recall and anti-polysaccharide responses^8^, yet failed to identify a universal signature of vaccination.

Here we leverage Immune Signatures Data Resource^26^, a curated database of publicly available datasets containing transcriptional and immune response profiling of peripheral blood following vaccination in humans, to perform a comparative analysis of transcriptional responses from 820 healthy young adults across 13 different vaccines. We find that while a common transcriptional program is shared across many vaccines, there is significant heterogeneity especially in the kinetics of immune responses. In particular, the live attenuated yellow fever vaccine induces a unique transcriptional response, with a surprisingly early upregulation of B and T cell modules within a day of vaccination, and a delayed induction of innate responses, including antiviral and interferon signaling, peaking at 10-14 days following vaccination. Furthermore, in an analysis of predictive signatures of antibody responses across vaccines, adjusting for time of peak expression enabled a gene module associated with plasma cells and immunoglobulins to consistently predict antibody responses across vaccines, demonstrating the importance of accounting for immune response kinetics in the development of universal predictors of response quality. Together, these findings highlight the spectrum of immune responses across vaccines and serve as a basis for future studies to understand the mechanisms underlying variation in immune responses across vaccines and inform future vaccine development.

## Results

### An integrated database of transcriptional responses to vaccination

As part of an effort to enable comparative studies and benchmarking of human vaccine responses, we curated a database of transcriptomic responses of 820 healthy adults (18-50 years old) across 13 different vaccines from previously published datasets. These datasets were compiled into *ImmPort*, an NIH-funded repository for immunological data^27^, and uploaded to *ImmuneSpace* (http://www.ImmuneSpace.org) for centralized QC and processing (Figure 1A). This combined database, named the Immune Signatures Data Resource^26^, includes responses to a broad range of vaccines, including live-attenuated viruses (e.g. yellow fever, smallpox and influenza vaccines), recombinant viral vectors (e.g. Ebola and HIV vaccines), inactivated viruses (e.g. seasonal influenza vaccine), glycoconjugate vaccines (e.g. pneumococcal and meningococcal vaccines) (Table S1). It also contains samples spanning multiple response timepoints, ranging from a few hours to more than 3 weeks post-vaccination (Figure 1B). Included participants in our initial analysis were restricted to 18-50 years old, and there were similar age and sex distributions across vaccines (Figure 1C). For analysis, all post-vaccination samples were normalized by pairwise fold change calculation with their matched pre-vaccination samples. Principal variance component analysis (PVCA)^28^ revealed that demographic features such as age and sex had relatively small contributions to variation in responses. In contrast, the post-vaccination timepoint of the sample explained 15% of the variance in the data, suggesting that there are shared kinetics of immune responses across vaccines (Figure 1D).

**Figure 1.**
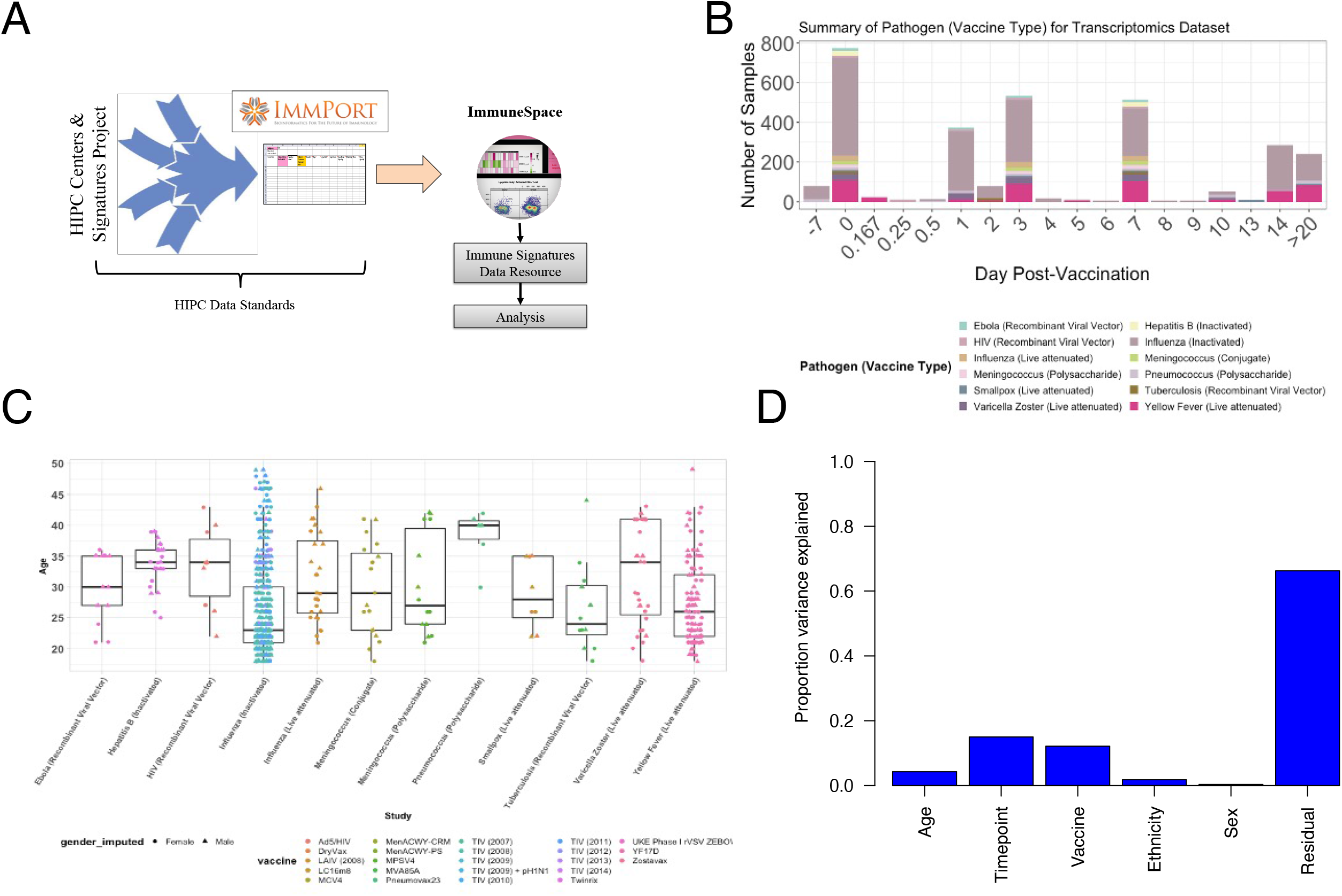
An integrated database of transcriptional responses to vaccination. A) Workflow for collection, curation, and standardization of datasets in the Immune Signatures Data Resource. B) Histogram of the number of samples included per vaccine at each timepoint in the Immune Signatures Data Resource. Day 0 represents Day of vaccination. C) Boxplots of the age distribution of participants in the Immune Signatures Data Resource by vaccine. Shape of points denotes the subject’s sex. D) Bar plot representing the proportion of variance in post-vaccination transcriptional responses that can be attributed to clinical (age, sex, ethnicity) and experimental variables (time after vaccination, vaccine) via Principal Component Variance Analysis. The residual represents the proportion of the variance that could not be explained by any of the included variables.

### Common and unique transcriptional responses across different vaccines

To examine the overlap in responses across vaccines, we identified differentially expressed genes post-vaccination relative to the pre-vaccination baseline as well as differential expression of blood transcriptional modules (BTMs), a set of gene modules developed through large-scale network integration of publicly available human blood transcriptomes^10^. There was much less overlap at a gene level (Figure S1A) than at a module level (Figure S1B), where a majority of differentially expressed modules were shared across 4 or more vaccines. Based on temporal expression patterns, the most commonly induced modules clustered into four groups (Figure 2A). Cluster 1 (indicated by the blue vertical bar to the left of the heatmap), upregulated at days 1 and 3 post-vaccination, represented BTMs related to innate responses and included modules associated with Toll-like receptor (TLR) and inflammatory signaling, antigen presentation, and monocyte signatures. Cluster 2 (yellow vertical bar to the left) contained multiple natural killer (NK) cell modules and was significantly downregulated on Day 1 (Figure S1C). Finally, Clusters 3 (pink bar) and 4 (green bar) generally peaked in activity on Day 7 and reflected plasma cell and cell cycle signatures, respectively, corresponding with expansion of antibody-producing plasmablasts. The “innate” Cluster 1 was most prominently induced in vaccines containing a live viral vector (Ebola, HIV), or an adjuvant (malaria) (Figure 2B). Meanwhile, the plasma cell signature in Cluster 3 was strongly increased in the polysaccharide pneumococcal vaccine and the conjugate meningococcal vaccine (Figure 2C).

**Figure 2.**
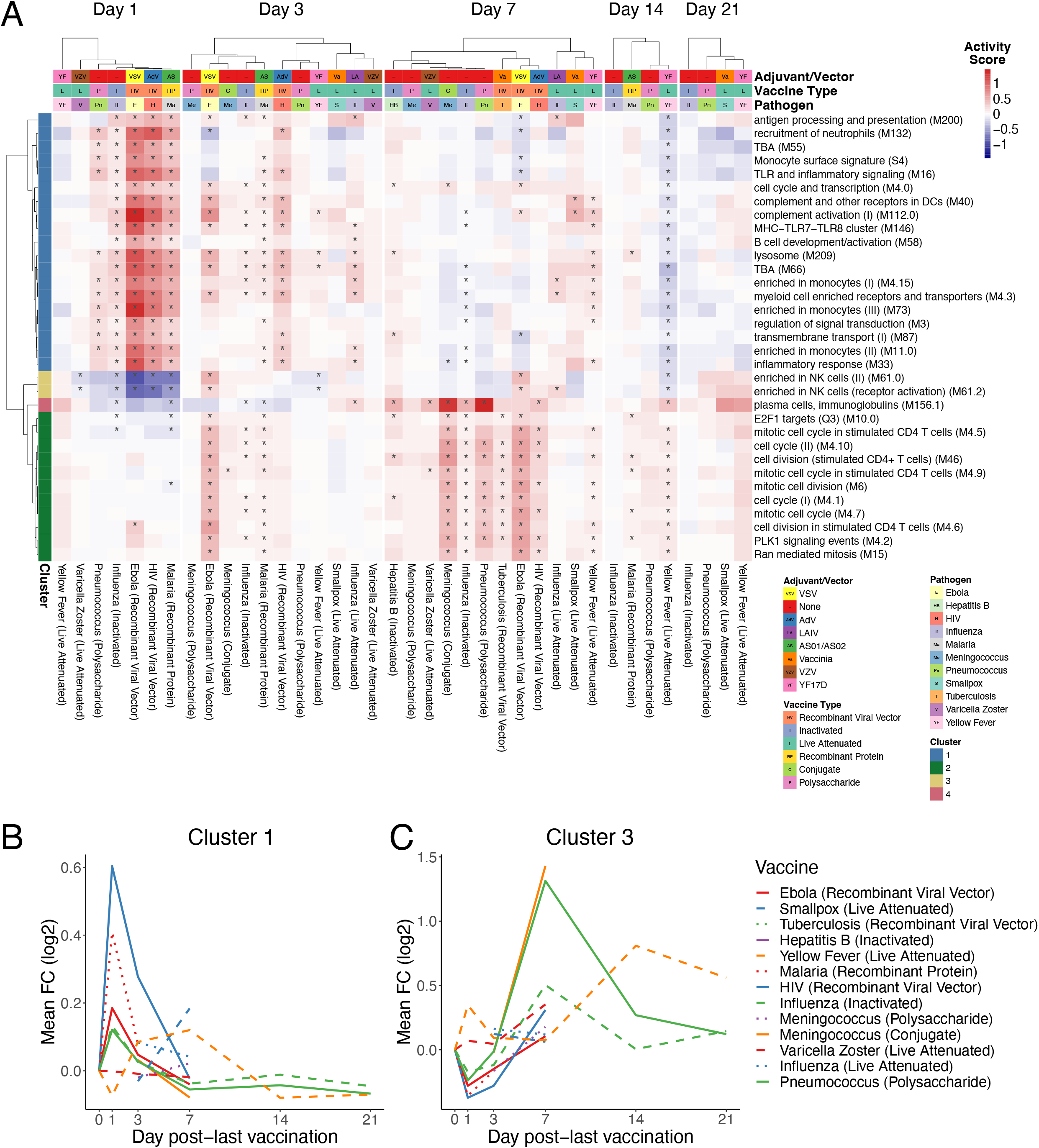
Common and unique transcriptional responses across different vaccines. A) Heatmap of common differentially expressed modules (regulated in 7 or more vaccines) over time (*FDR<0.05). Color represents the QuSAGE activity score. Clustering on columns was performed separately for Days 1, 3, 7, 14, and 21 post-vaccination. B) Kinetics of the mean FC of cluster 1 modules across vaccines C) Kinetics of the mean FC of cluster 3 modules across vaccines.

We next analyzed how differentially expressed modules were shared across vaccine responses (Figure 3). In agreement with the prior analysis, the response to most vaccines on Days 1 (Figure 3A) and 7 (Figure 3C) reflected innate and plasma cell/cell cycle responses, respectively, while the Day 3 response (Figure 3B) appeared as an intermediate between these states, with both innate and cell cycle signatures present. However, such responses were not universally shared across all of the vaccines. In particular, the early innate and antiviral responses common to most vaccines on Day 1 were not observed in the varicella zoster (VZV) and yellow fever vaccine responses. While these signatures appeared at later timepoints (Days 3 and 7) in yellow fever vaccine responses, they were not observed at all following VZV. Additionally, the Day 7 cell cycle signature was not observed following smallpox, VZV, and polysaccharide meningococcal vaccines. Notably, this signature was observed in the case of the meningococcal conjugate vaccine, where the bacterial polysaccharides have been conjugated to a diphtheria toxoid protein to induce memory and T helper cell responses^29^. Since diphtheria toxoid protein is used in other vaccines such as the Haemophilus influenza type B (Hib) vaccine^30^, the cell cycle signature observed at day 7 likely reflects the plasmablast response of the recall response to diphtheria toxoid, consistent with our previous study^10^.

**Figure 3.**
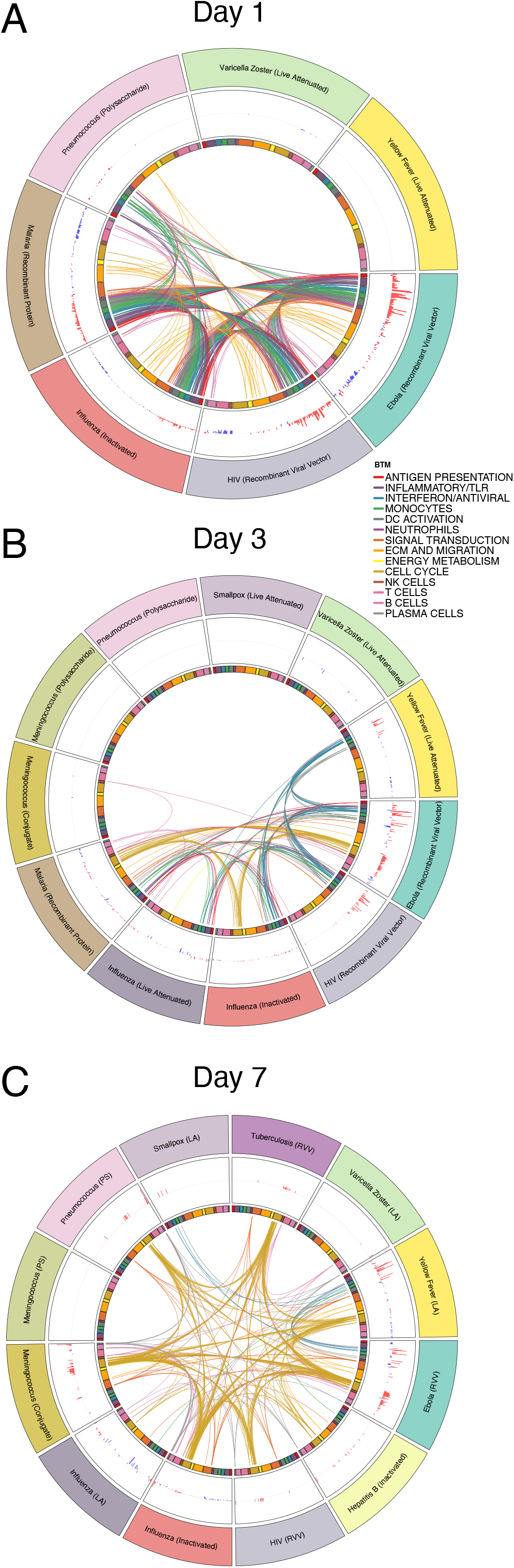
Overlap in transcriptional responses across vaccines. A-C) Circos plots of the overlap in differentially expressed modules (FDR<0.05) across vaccines on Days (A) 1, (B) 3, and (C) 7. Each segment of the circle represents one vaccine, and each point in a segment represents a single module. Bars in the outer circle represent the activity score of differentially expressed modules. Lines connect modules with a significant positive score shared between vaccines. Inner circle boxes and line colors represent the functional groups of the modules.

### Early adaptive and delayed innate transcriptional signatures of yellow fever vaccine

At the gene level, responses were highly correlated across most vaccines on Day 1 post-vaccination (Figure 4A) but became more divergent at later timepoints (Figures S2A-B). On Day 1, Ebola, inactivated influenza, HIV, and malaria vaccines exhibited the strongest similarity (Figures 4A, S2C). However, the yellow fever vaccine YF-17D induced a very distinct response that had little or even negative correlation with responses to all other vaccines, including other live viral vaccines such as VZV, HIV, and Ebola (Figures 4A, S2D). The innate pathways that were upregulated in other vaccine responses were, in fact, downregulated in response to yellow fever vaccine on Day 1 (Figure 4B). Instead, YF-17D induced early expression of multiple B and T cell modules. Analysis of estimated cell frequencies using the xCell deconvolution algorithm^31^ suggested that this induction may reflect a rapid increase in the frequency of peripheral B and CD4+ T cells (Figure S2E-F).

**Figure 4.**
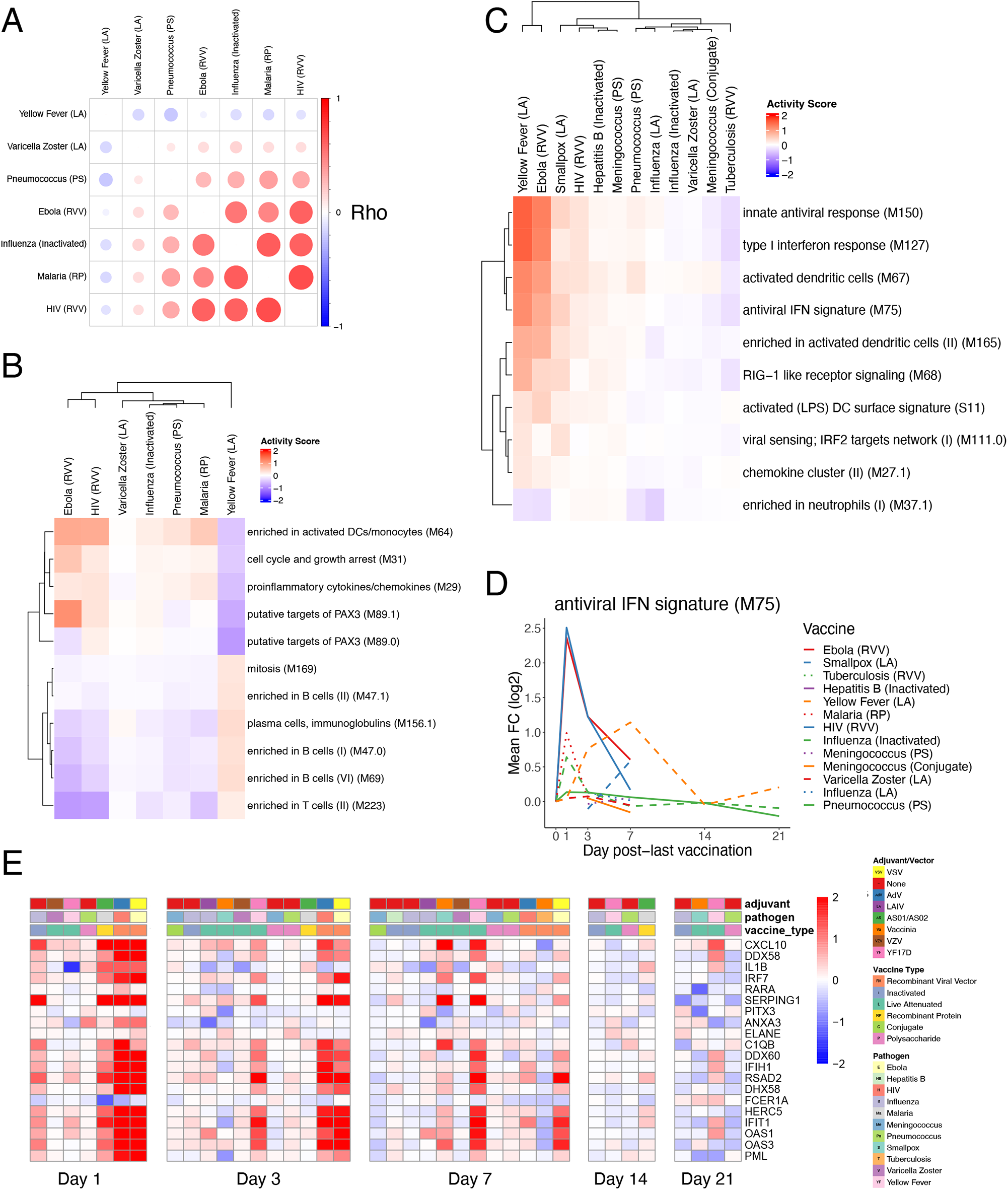
Early adaptive and delayed innate transcriptional signatures of yellow fever vaccine. A) Correlation matrix of pairwise Spearman correlations of Day 1 gene-level fold changes between vaccines. B) Heatmap of Day 1 activity scores of modules differentially expressed in response to YF vaccination (QuSAGE FDR<0.2). C) Heatmap of Day 7 activity scores of modules differentially expressed in response to YF vaccination (QuSAGE FDR<0.05, activity score >0.2). D) Kinetics of the mean FC of module M75 across vaccines. E) Heatmap of the post-vaccination FC of genes in module M75.

Another surprising feature of the yellow fever vaccine response was the relatively late expression of antiviral and interferon pathways, whose expression starts to be observed on Day 3 and peaks on Day 7 (Figure 4C-D). While these modules were also upregulated at this timepoint in Ebola vaccine responses, their expression waned rapidly following a robust early induction at Day 1. Some of the genes in these pathways that were strongly upregulated on Day 1 in response to most vaccines, such as CXCL10 and OAS1, were upregulated as late as 21 days post-vaccination with YF-17D (Figure 4E). Importantly, both the early adaptive and delayed innate responses were consistent across multiple studies from diverse geographical locations (Figures S2G-H). Together, these results highlight the unique kinetics of transcriptional responses to yellow fever vaccine relative to other vaccines.

### Time-adjusted transcriptional predictors of antibody responses

A key goal of systems vaccinology is to identify early signatures predictive of subsequent protection from infection. Antibody titers have been established as a reliable correlate of protection against many pathogens^32^ and previous studies have identified transcriptional signatures predictive of antibody responses to several vaccines, including inactivated influenza^6, 11, 18, 33, 34^, yellow fever^3^, and hepatitis B^12^. However, these signatures have thus far been developed for single vaccines, and it remains to be seen whether a ‘universal signature’ exists that can predict antibody responses across vaccines. Our curated data resource is uniquely suited to address this question, as 10 of the encompassed vaccines had at least one dataset with antibody titer measurements pre- and ∼1 month post-vaccination (Figure S3A). As there was substantial variability in antibody responses across vaccines, we defined ‘high’ and ‘low’ responders on a per dataset basis as the top and bottom 30% of participants according to antibody titer fold changes. We then used an elastic-net machine learning algorithm to develop classifiers capable of distinguishing between high and low responders based on early transcriptional signatures (see Methods section for further details).

As an initial approach, we wanted to examine whether a model trained using responses to a single vaccine could reliably predict responses to other vaccines. We therefore first built models using all 15 inactivated influenza vaccine datasets (the vaccine for which there was the largest number of samples) in a leave-one-study-out training/testing configuration. As validation that our model could predict responses within the same vaccine, classifiers trained using Day 7 fold-change expression data were able to predict high versus low antibody response in the left-out influenza dataset, with AUCs ranging between 0.55-0.9 (Figure 5A). The modules in these classifiers were primarily associated with cell cycle and plasma cell modules (Figure S3B). These results are consistent with prior work showing that classifiers built using similar pathways are predictive of antibody responses to influenza vaccination across multiple seasons^18^. However, when we examined their performance in other vaccines, they were not reliably predictive (Figure S3C). Moreover, the expression of modules associated with antibody response to inactivated influenza vaccination at Day 7 was not generally correlated with antibody responses across vaccines (Figure 5B).

**Figure 5.**
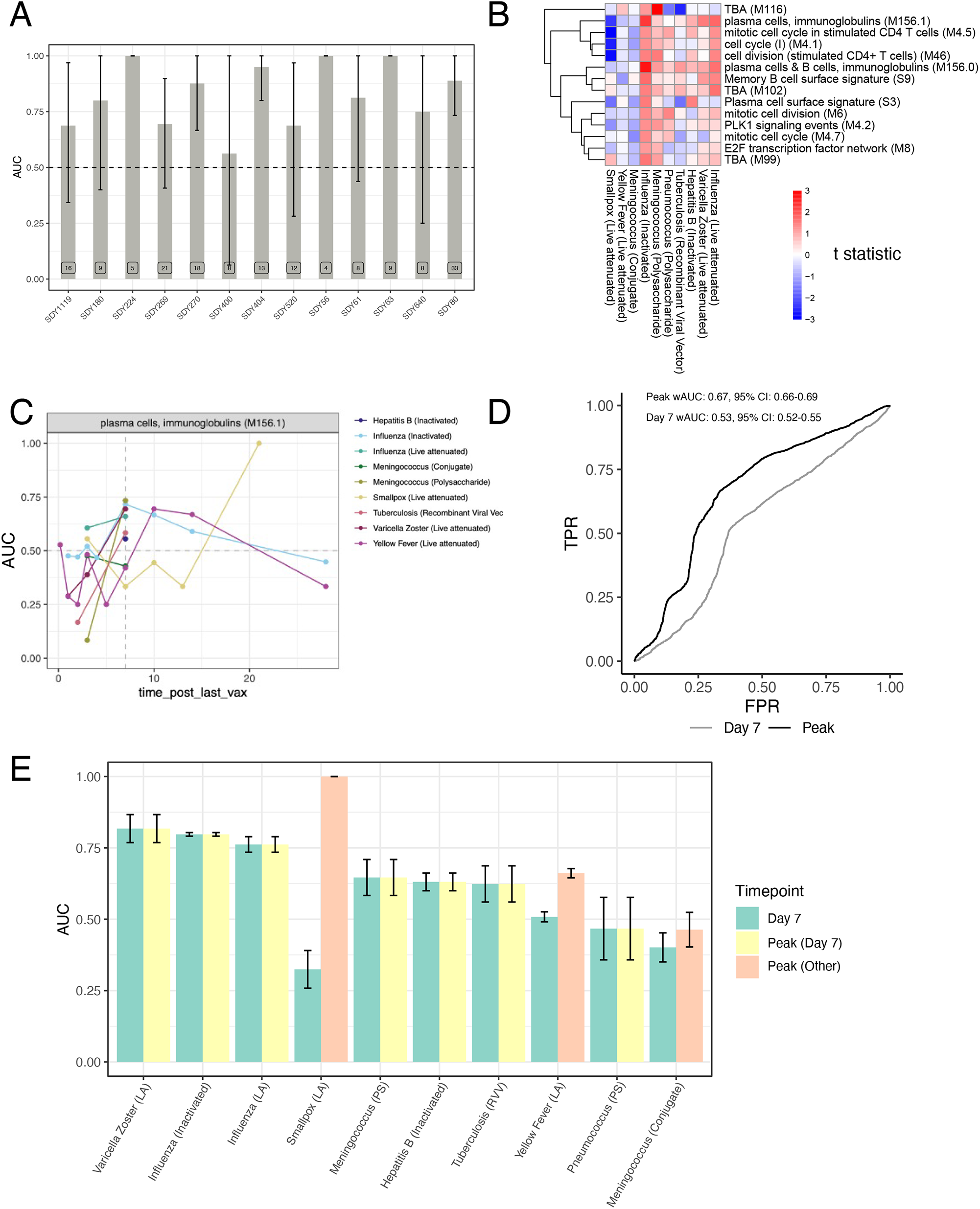
Time-adjusted transcriptional predictors of antibody responses. A) Area under the ROC curve (AUC) barplot of antibody response prediction performance per dataset for the elastic net classifier trained on inactivated influenza datasets only. B) Heatmap of high versus low antibody responder difference across vaccines of modules differentially expressed (FDR<0.05) between high and low antibody responders to inactivated influenza vaccination. C) Kinetics of the predictive power of M156.1 across vaccines. For each vaccine/timepoint combination, the AUC is computed based on difference in the geometric mean of the fold changes of the genes in the M156.1 between high and low responders (see Methods for details). D) Weighted ROC curves for a logistic regression classifier using M156.1 expression either at Day 7 in all vaccines (Day 7) or at the vaccine-specific peak expression timepoint (Peak) (Weighted AUC Day 7 / Peak = 0.65/0.53). E) Per vaccine AUC barplot for a logistic regression classifier using M156.1 expression either at Day 7 in all vaccines (yellow) or at the vaccine-specific peak expression timepoint (green – peak at Day 7, blue – peak at other timepoints).

We then asked whether training across multiple vaccines would improve the universality of the identified signatures. Neither a leave-one-vaccine-out approach, nor a 10-fold cross-validation approach combining all datasets, were able to identify signatures on Day 3 or Day 7 post-vaccination that could accurately discriminate high versus low responders across all vaccines (Figures S3D-E). However, analysis of the predictive power of specific modules over time, such as M156.1, one of the plasma cell modules associated with response in influenza vaccination on Day 7, revealed that this module was predictive of response across many vaccines but at different timepoints (Figure 5C). While many vaccines saw a strong association between M156.1 on Day 7 and subsequent antibody response, in certain vaccines such as yellow fever and smallpox, expression of the module was not associated with response until much later, at Days 10-14 and 21, respectively, consistent with the delayed kinetics of this BTM with these vaccines (Figure 2).

These results suggest that differential kinetics of immune responses across vaccines pose a confounding variable in the identification of universal predictive signatures of response at a single timepoint, but that using vaccine-specific timepoints dictated by the particular kinetics of immune responses for identification of predictive biomarkers of vaccine responses may improve the universality of such signatures. To test this hypothesis in the context of the plasma cell signature, we identified the timepoint at which expression of the plasma cell module M156.1 peaked in response to each vaccine (Figure 2C). We then trained a logistic regression classifier with M156.1 expression as an input in a 10-fold cross-validation approach using fold-change data at the peak M156.1 expression timepoint for each vaccine. Indeed, using M156.1 peak expression timepoints improved the overall performance of the classifier compared with using a single timepoint (Day 7) for all of the vaccines (Figure 5D). This improvement was driven by increases in response prediction among vaccines in which the plasma cell signature peaked at timepoints other than Day 7, such as the yellow fever and smallpox vaccines (Figure 5E). Thus, expression of the plasma cell module M156.1 acts as a time-variable universal signature of antibody responses to vaccination.

### Impact of aging on transcriptional responses to vaccination

The impairment of vaccine efficacy with age is a major challenge for vaccine development and public health. Although declining vaccine efficacy can broadly be attributed to effects of immunosenescence such as loss and dysfunction of naïve T cells^35^, diminished class-switch capability of B cells^36^, and decreased TLR function among innate cells^37, 38^, the molecular mechanisms responsible for impaired vaccine responses among older adults are not yet fully understood. While most of the curated datasets in the HIPC resource contained only young adult participants, some studies, including those of inactivated influenza^18^, varicella zoster^13^, and hepatitis B^12^ vaccines, profiled responses of both young (≤50) and older (≥60) vaccinees. As expected, post-vaccination antibody responses were diminished in older compared to younger participants across all three vaccines (Figure 6A).

**Figure 6.**
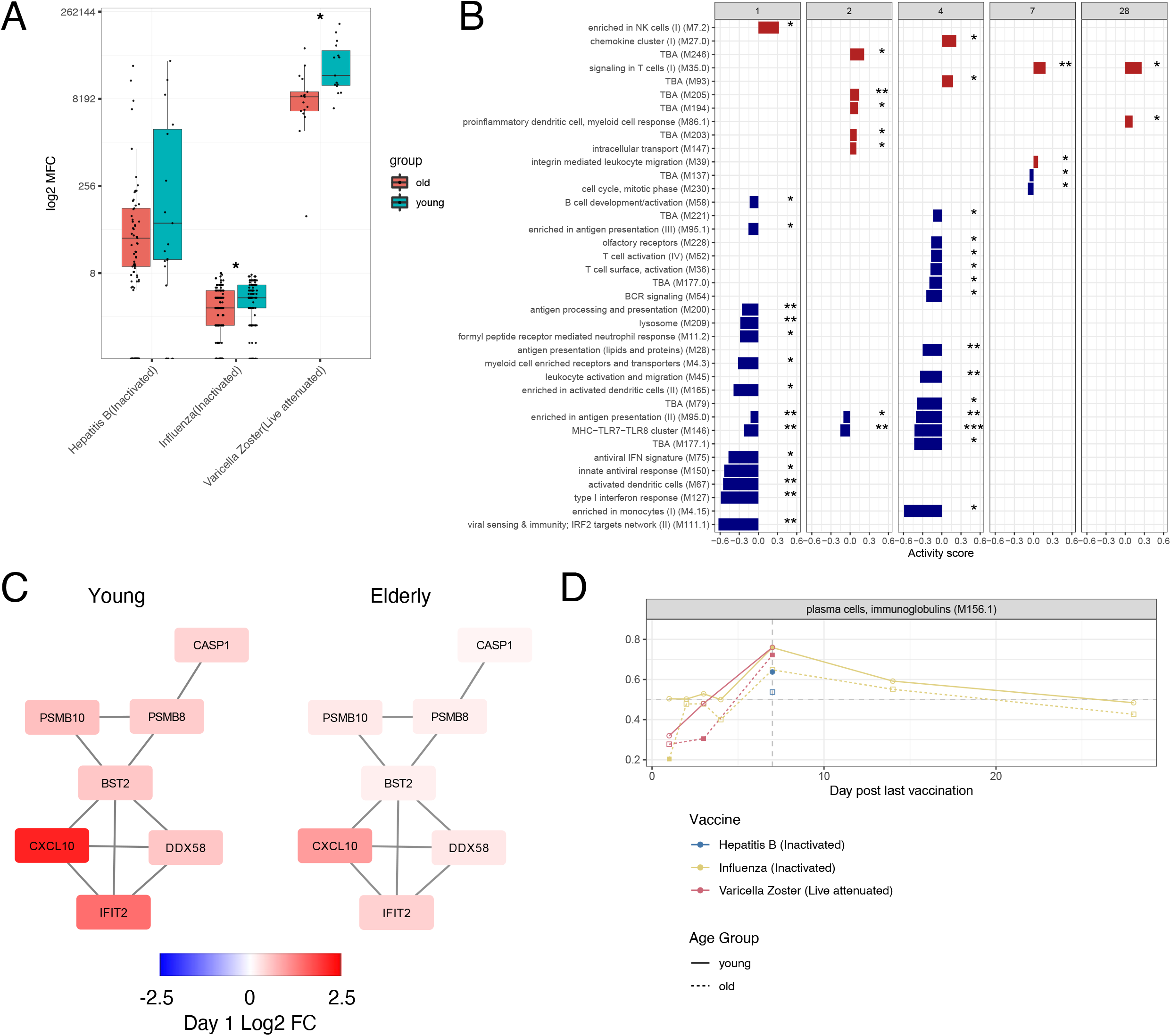
Impact of aging on transcriptional responses to vaccination. A) Boxplots of Day 30 antibody responses to vaccination in young (≤50) and older (≥60) participants across vaccines. B) Modules differentially expressed between young and older participants in response to inactivated influenza vaccination (QuSAGE FDR<0.05). C) Network plot of modules M111.1 on Day 1 following inactivated influenza vaccination in young and older participants. Each edge represents a co-expression relationship, as described in Li et al.^10^; colors represent the Day 1 log2 FC. D) Kinetics of the predictive power of modules M156.1 across vaccines in young and older participants (filled circles indicate p<0.05, 1000 permutations).

We sought to examine for the effect of aging on immune responses across vaccines by comparing BTM activity scores of the most commonly induced BTMs (Figure 2A) between young and older participants across all three vaccines at each timepoint. Broadly, transcriptional responses to the three vaccines were similar between the two age groups (Figure S4A). However, there were significant age-associated differences in several pathways in response to inactivated influenza vaccination, including decreased expression of interferon and other innate immune modules in older compared to young participants early post-vaccination (Figure 6B-C), consistent with prior findings^18^. Despite these differences, the power of the M156.1 plasma cell module to predict the antibody response was highly similar in both young and older individuals (Figure 6D). These results suggest conservation in the pathways responsible for successful antibody production post-vaccination, consistent with prior findings for influenza vaccination^20^.

## Discussion

The high degree of homology in the vaccine-induced signatures induced demonstrates that diverse vaccines that differ widely in target pathogens and composition stimulate conserved immunological networks. Despite this homology, there was still substantial heterogeneity in both the magnitude and kinetics of the induced responses across vaccines. The most distinct in this regard were responses to the yellow fever vaccine YF-17D, which displayed several unique features: (1) a delayed innate and antiviral response which did not peak until Days 3-7 post-vaccination (Figure 4D), (2) an early upregulation of B and T cell signatures at Day 1 (Figures 4B, S3E) not observed in other vaccines until much later, and (3) a delay in cell cycle and plasma cell signatures typically associated with the expansion of antigen-specific antibody-secreting cells (Figures 2A-B).

The mechanisms underlying the delayed responses are unclear, but could be caused by differences in viral tropism, the slow but sustained tempo of viral replication *in vivo*, or unique immune-evading properties of YF-17D. Wild-type yellow fever infects both Kupffer cells and hepatocytes in the liver, with the potential for severe pathology^39^; however data from NHPs suggests that YF-17D infects lymphoid cells at the site of injection and spreads to monocytes and macrophages in the lymph nodes, bone marrow, and spleen but does not infect the liver^40, 41^. This tropism appears similar to other live viral or viral-vectored intramuscular vaccines included in this dataset such as rVSV-ZEBOV which is thought to target endothelial cells, monocytes, macrophages, and myeloid dendritic cells in lymphoreticular tissues^42^, and MRKAd5/HIV containing an Ad5 adenovirus vector, which also has broad tropism but appears to cause local and lymphoreticular infection without reaching the liver following intramuscular administration^43, 44^.

Of note, yellow fever and other flaviviruses have a specific capability to inhibit interferon signaling via multiple mechanisms, including suppression of JAK-STAT signaling^45^, which could potentially cause the observed delay in interferon responses following YF-17D vaccination. Interestingly, the *Vaccinia* virus also has several mechanisms for inhibition of interferon responses, including prevention of IRF-3 and NFκB activation and dephosphorylation of STAT1/2^46^. Although early response data was not available, the smallpox vaccine containing *Vaccinia* also induced some degree of delayed interferon response following vaccination (Figure 4D).

While YF-17D demonstrated delayed induction of interferon signatures, induction of B and T cell signatures at Day 1 was much earlier than typically observed with other vaccines. This timing is most likely too early to represent an antigen-specific response but could reflect non-specific activation or recruitment of naïve cells into the circulation. Alternatively, these signatures could be a result of increased relative proportions of adaptive cells in the blood due to extravasation of innate cells into tissues at the site of injection. Further investigation at a cellular level is required to address these hypotheses and elucidate the mechanisms by which YF-17D exerts such unique early effects on the adaptive immune system.

Finally, our analysis of predictive signatures of antibody responses (Figure 5) indicates that vaccine response kinetics play an important role in determining such signatures. Here we have illustrated this principal for a single plasma cell transcriptional module, however future analyses may enable detection of additional and more accurate signatures. We have previously proposed the concept of a ‘vaccine chip’ that could measure defined biomarkers and be used to predict protective immune responses across vaccines^25^. This chip would be designed to measure expression of a select set of genes or modules, subsets of which would predict a particular type of functional or protective immune response (e.g., neutralizing antibody titers, effector CD8+ T cell responses, frequency of polyfunctional T cells, T helper 1 (T_h_1) versus T_h_2 response bias, etc). Our findings demonstrate that the unique kinetics of immune responses to different vaccines should be accounted for in the development of such a tool. In practice, small phase I/II trials could be used to define response kinetics and enable the successful application of a ‘vaccine chip’ to predict immune responses in subsequent trials.

Due to the significant costs needed to perform a clinical trial of sufficient size, such vaccine studies are rarely performed with more than one vaccine. Here, we have demonstrated that meta-analysis of vaccine trials can provide valuable insights into the common and unique aspects of immune responses across vaccines. Combined with the Immune Signatures Data Resource^26^, these computational approaches and repositories will enable future research into the mechanisms of vaccine-induced immunity to inform development of improved adjuvants and vaccines.

## Acknowledgements

This research was performed as a project of the Human Immunology Project Consortium (HIPC) and supported by the National Institute of Allergy and Infectious Diseases. This work was supported in part by NIH grants U19AI118608, U19AI128949, U19AI090023, U19AI118626, U19AI089992, U19AI128914, U19AI128910, U19AI118610, U19AI128913, and the Intramural Program of NIAID and NIH institutes supporting the Trans-NIH Center for Human Immunology (CHI). OL is supported in part by the Department of Pediatrics at Boston Children’s Hospital. Work in Bali Pulendran’s lab is supported in part by National Institutes of Health (R37 DK057665; R01 AI048638; U19 AI057266; U19 AI167903), Bill and Melinda Gates Foundation, Open Philanthropy, and the Violetta L. Horton and Soffer Endowments to B.P.

## Methods

### Gene expression preprocessing

An extensive description of the preprocessing of microarray and RNA-Sequencing (RNA-Seq) datasets included in the Immune Signatures Data Resource can be found in the associated manuscript^26^. The dataset includes 2,949 samples from published studies and 228 samples not included in previously published studies. All these samples were assembled into a single resource. Briefly, raw probe intensity data for Affymetrix studies were background corrected and summarized using the RMA algorithm^47^. For studies using the Illumina array platform, background corrected raw probe intensities were used. For RNA-Seq studies, count data was voom-transformed^48^ to mimic the distribution of microarray expression intensities. Expression data within each study was quantile normalized and log-transformed separately for each study.

### Batch correction

An extensive description of the across studies normalization used to correct for batch effects can be found in the Immune Signatures Data Resource manuscript^26^. Briefly, a linear model was fit using the pre-vaccination normalized gene expression as a dependent variable and platform, study, and blood sample type (i.e., whole blood or PBMC) as independent variables. The estimated effect of the platform, study and sample type was then subtracted from the entire gene expression (pre- and post-vaccination) to obtain batch corrected gene expression.

### Identification of differentially expressed genes

To determine differentially expressed genes, p values were first computed within each study using paired student’s t-tests. Next, Stouffer’s method was used to combine p values across studies via the *sumz* function in the metap R package^49^, with weighting according to the square root of the study sample size. Finally, combined p values were then adjusted for multiple testing using the Benjamini-Hochberg procedure. Similarly, average gene fold changes for each vaccine at each timepoint were computed by averaging across studies while using weighting equal to the study sample size.

### Gene set enrichment analysis

The enrichment analysis of BTMs was performed in two steps. First, for every study and time point, enrichment was calculated using QuSAGE^50^, providing as contrast “Day X – Day 0” where X is the current time point, and also a “pairVector” containing the subject identifiers so that a paired analysis would be performed. Second, to integrate the results from multiple studies of the same vaccine, we performed a meta-analysis for every vaccine + timepoint combination, using the “combinePDFs” function of QuSAGE.

### Gene and module sharing analysis

The sharing number of a gene/module is computed as the maximum number of vaccines it is significantly differentially expressed (FDR<0.05) in, irrespective of time point. For modules, the p-values were calculated using QuSAGE^50^ (see *Gene set enrichment analysis*). A null distribution for sharing was generated by performing 10,000 permutations of gene/module labels within each vaccine + timepoint group.

### Antibody titer measurements and identification of high and low responders

Depending on the study, antibody titers were measured by neutralization assays, hemagglutination inhibition assay (HAI), or Immunoglobulin G (IgG) levels measured by ELISA^26^. Since some vaccines include multiple strains of viral antigens, the fold change in the antibody response metric was defined as the maximum fold change (MFC) of any strain in the vaccine at day 28 (+/-7 days) compared to pre-vaccination. To minimize the difference in antibody response between studies (e.g., due to different vaccines or different techniques used for antibody concentration assessment), the high and low responders were identified for each study separately by selecting the participants with MFC equal or above the 70th percentile as high responders and participants with MFC equal or below the 30th percentile as low responders.

### Identification of predictive signatures of antibody responses

Four training/testing setups employed for identification of predictive signatures of antibody responses: 1) inactivated influenza datasets only, leave-one-study-out 2) training on all inactivated influenza datasets, testing on other vaccines 3) leave-one-vaccine-out (all datasets combined) 4) 10-fold cross validation (all datasets combined). All models were trained using elastic-net logistic regression using the ‘caret’ and ‘glmnet’ R packages. BTM enrichment scores were calculated for each sample using the single-sample Gene Set Enrichment Analysis (ssGSEA) function and used as input features to the models, filtering for modules with a standard deviation > 75% quantile of the standard deviation. Models were fit using either Day 3 fold-change or Day 7 fold-change of ssGSEA score separately. Tuning parameters and performance metrics were estimated using 10-fold cross-validation. Confidence intervals were estimated using the ‘ci.auc’ function from the pROC R package.

When developing predictive models for the timepoint adjustment approach, logistic regression models were trained using ssGSEA score fold-change for module M156.1, either at Day 7 or at the timepoint of peak expression in a given vaccine. AUC confidence intervals were estimated using linear-mixed effects models fitted with 100 Monte-Carlo resamples. When computing AUCs across multiple vaccines, a weighted AUC was computed using sample size as the weights. For the analysis of temporal change in the predictive capability of M156.1 (Figure 4C), a weighted mean AUC (based on number of samples in each study) was computed using the calculateROC function of MetaIntegrator R package based on the geometric mean of gene fold changes in the M156.1 module.

## Figure Captions

**Table S1.** Summary of vaccine datasets.

| ImmPort Accession | Pathogen | Vaccine Type | Adjuvant/Vector | Sample Type | # of samples |
| --- | --- | --- | --- | --- | --- |
| SDY1373 | Ebola | Recombinant Viral Vector | VSV | Whole blood | 46 |
| SDY1328 | Hepatitis B | Inactivated | None | Whole blood | 51 |
| SDY1291 | HIV | Recombinant Viral Vector | AdV | PBMC | 50 |
| SDY1119 | Influenza | Inactivated | None | PBMC | 67 |
| SDY1276 | Influenza | Inactivated | None | Whole blood | 828 |
| SDY180 | Influenza | Inactivated | None | Whole blood | 102 |
| SDY212 | Influenza | Inactivated | None | Whole blood | 29 |
| SDY224 | Influenza | Inactivated | None | PBMC | 55 |
| SDY269 | Influenza | Inactivated | None | PBMC | 80 |
| SDY270 | Influenza | Inactivated | None | PBMC | 83 |
| SDY400 | Influenza | Inactivated | None | PBMC | 60 |
| SDY404 | Influenza | Inactivated | None | PBMC | 64 |
| SDY520 | Influenza | Inactivated | None | Whole blood | 51 |
| SDY56 | Influenza | Inactivated | None | PBMC | 96 |
| SDY61 | Influenza | Inactivated | None | PBMC | 27 |
| SDY63 | Influenza | Inactivated | None | PBMC | 42 |
| SDY640 | Influenza | Inactivated | None | Whole blood | 44 |
| SDY80 | Influenza | Inactivated | None | PBMC | 256 |
| SDY269 | Influenza | Live attenuated | LAIV | PBMC | 83 |
| SDY1293 | Malaria | Recombinant protein | AS01/AS02 | PBMC | 165 |
| SDY1260 | Meningococcus | Conjugate | None | PBMC | 51 |
| SDY1325 | Meningococcus | Conjugate | None | Whole blood | 4 |
| SDY1260 | Meningococcus | Polysaccharide | None | PBMC | 39 |
| SDY1325 | Meningococcus | Polysaccharide | None | Whole blood | 2 |
| SDY180 | Pneumococcus | Polysaccharide | None | Whole blood | 54 |
| SDY180 | Pneumococcus | Polysaccharide | None | Whole blood | 101 |
| SDY1370 | Smallpox | Live attenuated | Vaccinia | PBMC | 48 |
| SDY1364 | Tuberculosis | Recombinant Viral Vector | Vaccinia | PBMC | 36 |
| SDY984 | Varicella Zoster | Live attenuated | VZV | PBMC | 124 |
| SDY1264 | Yellow Fever | Live attenuated | YF17D | PBMC | 87 |
| SDY1289 | Yellow Fever | Live attenuated | YF17D | Whole blood | 117 |
| SDY1294 | Yellow Fever | Live attenuated | YF17D | PBMC | 109 |
| SDY1529 | Yellow Fever | Live attenuated | YF17D | Whole blood | 180 |

**Figure S1.**
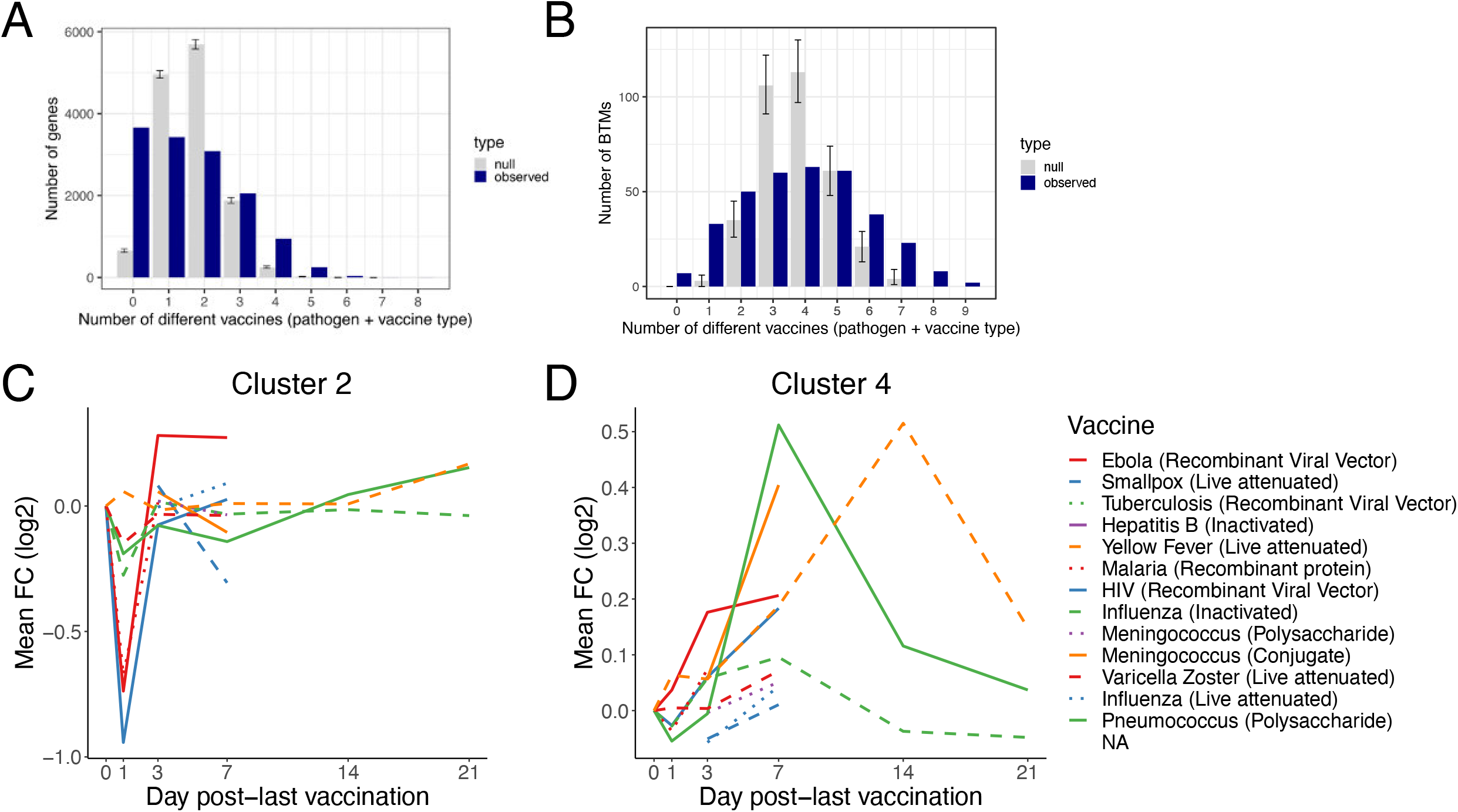
Overlap in differentially expressed genes/modules and kinetics of common module clusters. A-B) Histograms of overlap in DEGs (A) or differentially expressed modules (B) between vaccines. A gene/module is shared with another vaccine if it is significantly (FDR < 0.05) regulated in the same direction, irrespective of time point. Grey bars represent the null distribution generated by 10,000 permutations of gene/module labels within vaccine + timepoint groups. Error bars indicate the 2.5% and 97.5% quantiles. C) Kinetics of the mean FC of cluster 2 BTMs across vaccines. D) Kinetics of the mean FC of cluster 4 modules across vaccines. A gene(/set) is shared with another vaccine if it is significantly (FDR < 0.05) up/down in the same direction, irrespective of time point. Blue bars, number of genesets shared (y-axis) between the same number of vaccines (x-axis). Grey bars, null distribution generated by 10k permutations of geneset labels within vaccine+timepoint groups. Error bars indicate the 2.5% and 97.5% quantiles.

**Figure S2.**
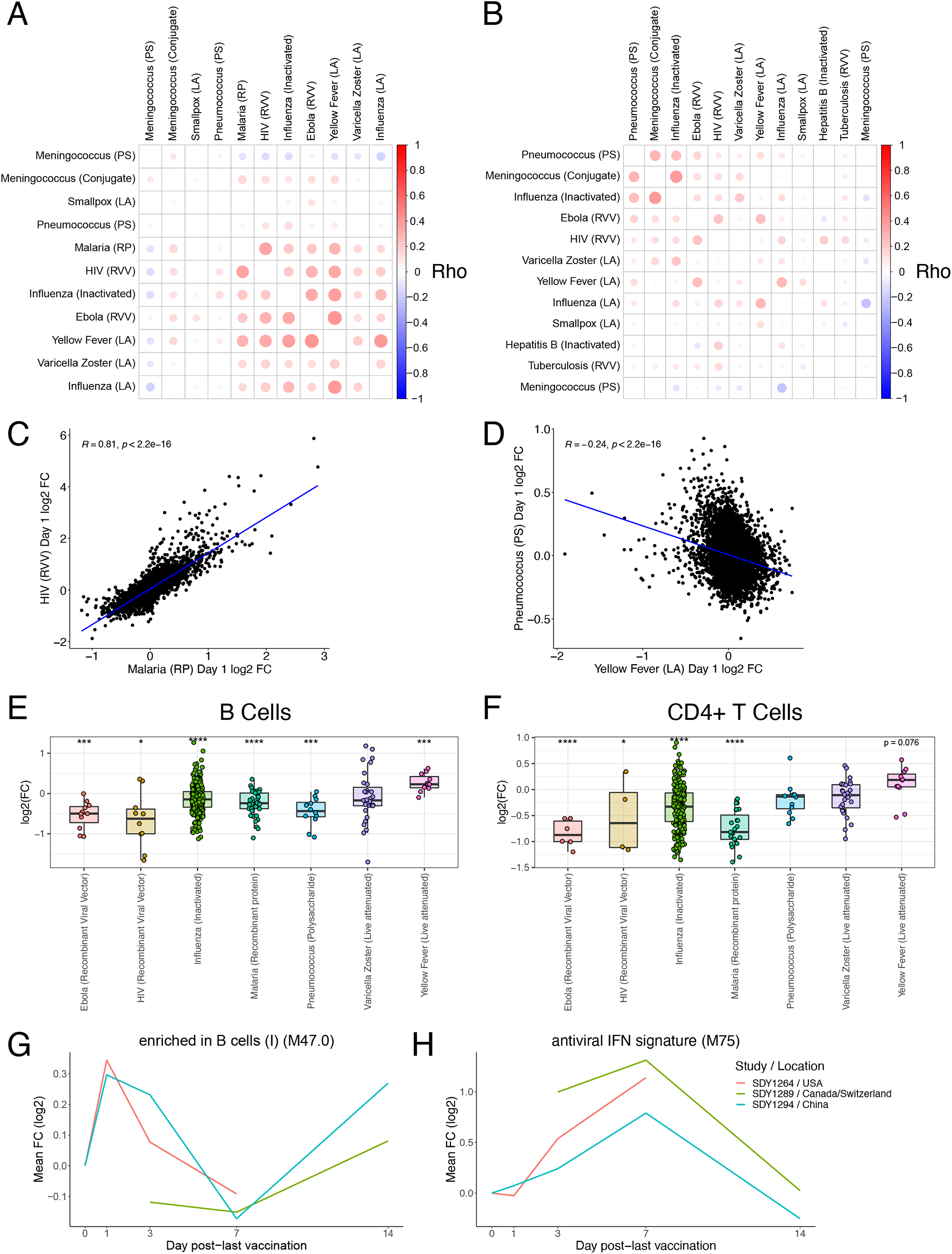
Gene-level correlations between vaccines and estimated cell frequencies. A) Correlation matrix of pairwise Spearman correlations of Day 3 gene-level fold changes between vaccines. B) Correlation matrix of pairwise Spearman correlations of Day 7 gene-level fold changes between vaccines. C) Scatterplot of Day 1 gene FCs between HIV and Malaria vaccines. D) Scatterplot of Day 1 gene FCs between Yellow Fever and Pneumococcus vaccines. E) Boxplot of Day 1 FC in xCell^31^ estimated B cell frequencies across vaccines. F) Boxplot of Day 1 FC in xCell^31^ estimated CD4+ T cell frequencies across vaccines. *p < 0.05, **p < 0.01, ***p < 0.001, **** p < 0.0001.

**Figure S3.**
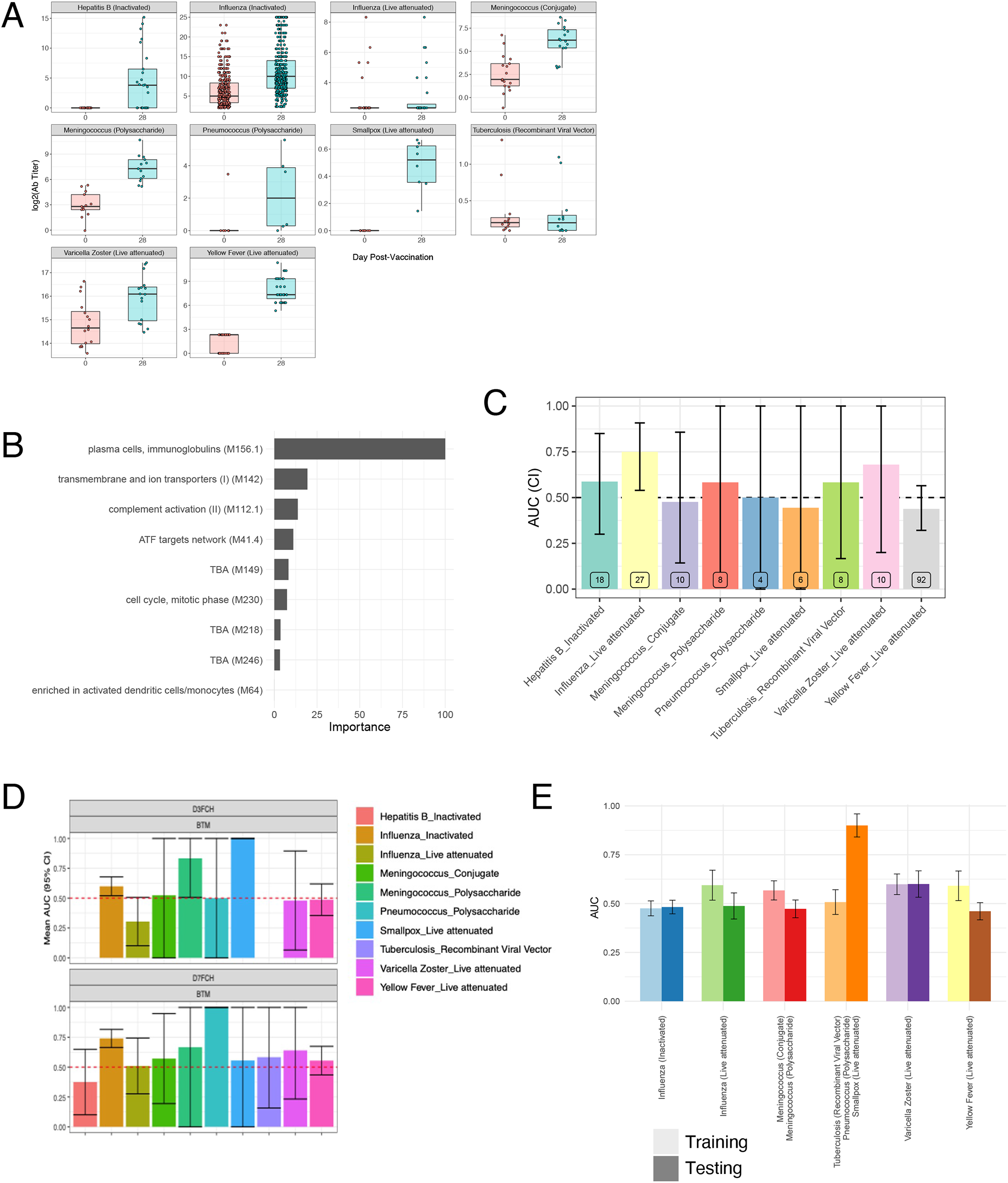
Antibody response prediction across vaccines. A) Boxplots of Day 30 antibody responses to vaccination across vaccines. B) Barplot of feature importance for the GLM classifier trained on inactivated influenza datasets only. AUC barplot of antibody response prediction performance across vaccines for the GLM classifier trained on inactivated influenza datasets only. C) AUC barplot of antibody response prediction performance of the leave-one-vaccine-out GLM classifier. D) AUC barplot of antibody response prediction performance of the 10-fold cross-validation GLM classifier.

**Figure S4.**
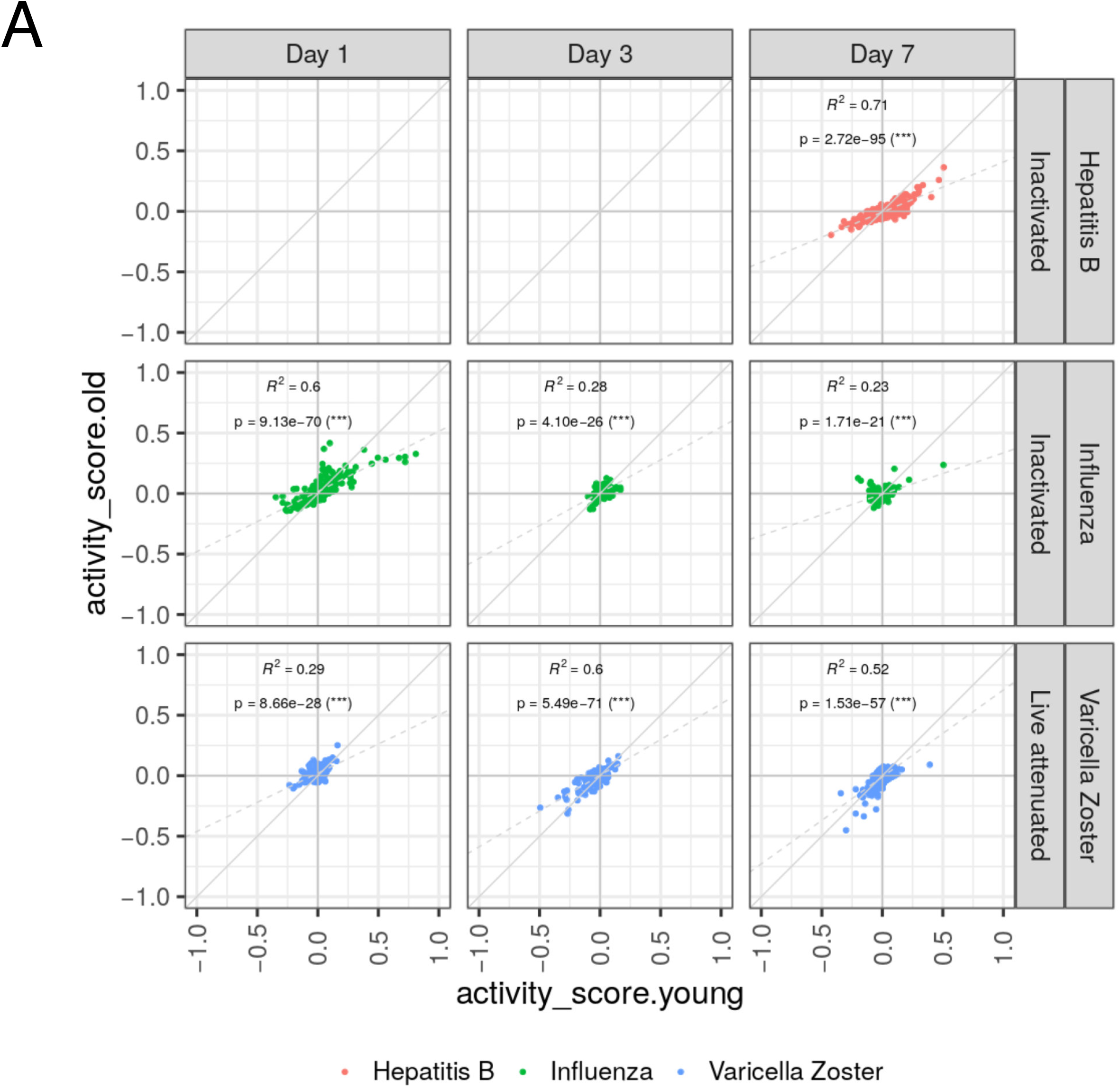
Comparison of common transcriptional responses between age groups. A) Scatterplots of module activity scores in each vaccine among young (x-axis) and elderly (y-axis) of the most commonly expressed modules (Figure 2A) on days 1-7.

